# An exploration of assembly strategies and quality metrics on the accuracy of the *Knightia excelsa* (rewarewa) genome

**DOI:** 10.1101/2020.10.28.358903

**Authors:** Ann McCartney, Elena Hilario, Seung-Sub Choi, Joseph Guhlin, Jessica M. Prebble, Gary Houliston, Thomas R. Buckley, David Chagné

## Abstract

**Background:** We used long read sequencing data generated from *Knightia excelsaI* R.Br, a nectar producing Proteaceae tree endemic to Aotearoa New Zealand, to explore how sequencing data type, volume and workflows can impact final assembly accuracy and chromosome construction. Establishing a high-quality genome for this species has specific cultural importance to Māori, the indigenous people, as well as commercial importance to honey producers in Aotearoa New Zealand.

**Results:** Assemblies were produced by five long read assemblers using data subsampled based on read lengths, two polishing strategies, and two Hi-C mapping methods. Our results from subsampling the data by read length showed that each assembler tested performed differently depending on the coverage and the read length of the data. Assemblies that used longer read lengths (>30 kb) and lower coverage were the most contiguous, kmer and gene complete. The final genome assembly was constructed into pseudochromosomes using all available data assembled with FLYE, polished using Racon/Medaka/Pilon combined, scaffolded using SALSA2 and AllHiC, curated using Juicebox, and validated by synteny with *Macadamia*.

**Conclusions:** We highlighted the importance of developing assembly workflows based on the volume and type of sequencing data and establishing a set of robust quality metrics for generating high quality assemblies. Scaffolding analyses highlighted that problems found in the initial assemblies could not be resolved accurately by utilizing Hi-C data and that scaffolded assemblies were more accurate when the underlying contig assembly was of higher accuracy. These findings provide insight into what is required for future high-quality *de-novo* assemblies of non-model organisms.

## Background

It is of critical importance that an optimal genome assembly strategy is used to maximize the impact, effectiveness, and accuracy of resulting pseudo-chromosome-scale *de novo* reference genomes. As long read sequencing data becomes more affordable, the integration of a multitude of next generation sequencing (NGS) platforms is becoming standard for generating near-complete *de novo* genome assemblies. The construction of an accurate *de novo* assembly is crucial to facilitating investigations of species evolution (“https://vertebrategenomesproject.org/,”; “https://www.darwintreeoflife.org/,”), organism diversity (Gurdasani D., Martinez J., Pollard M., Carstensen T., & C., 2016; “https://www.nist.gov/programs-projects/genome-bottle,”; Project), and informing health and disease treatments in fields such as cancer treatment programs (Berger & Mardis, 2018) and vaccine development (Prachi, Donati, Masciopinto, Rappuoli, & Bagnoli, 2013). To cater for the synergistic nature of different types of sequencing data the research field of genome assembly is moving quickly, and new methods are becoming more flexible, accurate and efficient. Genome assembly software incorporates sophisticated algorithms built to deal with a multitude of sequencing data types; for instance, accounting for the different base calling accuracies of Oxford Nanopore Technology (ONT) (<5% error rate), PacBio Single Molecule Real-Time (SMRT) (<1% error rate) and Illumina short paired-end (PE) reads (<0.1% error rate). They also allow a multitude of parameter specifications to cater for various genome architectures. For example, centroFlye (Bzikadze & Pevzner, 2019) is designed for centromere assembly, and chloroExtractor (Ankenbrand et al., 2018) developed to assemble chloroplastic genomes from whole genome sequencing (WGS) data. Through different error correction and consensus approaches, these software use noisy ONT data to construct contig assemblies which can then be further scaffolded to generate high-quality assemblies, but it is not generally clear what type of data are required or the volume necessary to generate the ‘optimal’ assembly, or indeed what combination of software one should use given the available types and volumes of data.

Despite a thorough investigation of the computational resource performance of long read assemblers in 201 9(Wick & Holt, 2019), published data on the optimization of read length and depth in the context of the most commonly used long read assemblers (NECAT, WTDBG2/RedBean, CANU, FLYE, FALCON and SHASTA) is limited. Although the underlying long read data used by these toolkits were shared, their methods for error correction, assembly and consensus generation differ greatly. For instance, FLYE (Kolmogorov, Yuan, Lin, & Pevzner, 2019) identifies “disjointigs” and uses these to firstly resolve the repeat graph in order to construct the final assembly. CANU (Koren et al., 2017) carries out extensive error correction and trimming prior to generating the final assembly using overlap-consensus methods based on string graph theory (Myers, 2005). NECAT (Chen et al., 2020) acts similarly to CANU albeit using a more progressive correction and assembly strategy. In contrast, WTDBG2/RedBean (Ruan & Li, 2020) uses only a single round of consensus by a fuzzy DeBruijn algorithm (Zerbino & Birney, 2008) that is based on initial short read assembly algorithms that have been adjusted to accommodate the base calling inaccuracies of noisy long reads. The SHASTA (Kishwar Shafin et al., 2020) algorithm maximizes computational efficiency through the identification of reduced marker kmers to initially find overlaps and then build the consensus sequence.

Gaining an understanding of each assembler’s advantages and shortcomings is an important consideration prior to assembly to form a more educated assembly strategy and ultimately resulting in a genome assembly sufficient for individual project needs. Quantitative metrics to track the accuracy and completeness of the assembly must be performed as often as possible throughout the workflow. In the past, appropriate non-manual methods of genome accuracy assessment have been limited, particularly in regards to scaffolding steps using as proximity-guided methods like Hi-C (Lieberman-Aiden et al., 2009). Recently, more advanced quantitative toolkits have become available, such as kmer completeness [Merqury (Rhie, Walenz, Koren, & Phillippy, 2020)], Long terminal repeat retrotransposons Assembly Index [LAI (Ou, Chen, & Jiang, 2018)], mapping rate and highly conserved gene completeness [BUSCO (Simão, Waterhouse, Ioannidis, Kriventseva, & Zdobnov, 2015)]. However, an educated selection of the assembler only is insufficient in generating an optimal genome assembly, as post processing steps such as polishing and scaffolding are also important considerations.

The identification and correction of mis-assemblies, or “polishing,” is determined by the initial assembler and the algorithm used, however, comprehensive analyses of the impact of different polishing strategies on genome accuracy are scarce. Assembler algorithms act differently during contig construction; thus, the initial assembly accuracy they produce before polishing is not always a fair indication of the metrics that will be obtained afterwards. Iterative polishing steps increase assembly accuracy after each step so that reads previously unable to map due to error or mis-assembly in the initial assembly become mappable, indicating a more accurate consensus assembly. Polishers are placed in two categories “Sequencer bound” or “General”. Both Nanopolish (Loman, Quick, & Simpson, 2015) and Medaka (Technologies, 2018) are examples of sequencer bound polishers that utilize raw signal information while Racon (Vaser, Sović, Nagarajan, & Šikić, 2017) and Pilon (Walker et al., 2014) are examples of general polishers that are applicable to any sequencing platform. To obtain a better understanding of polishing and post assembly processing performance, initial contig assemblies generated from a selection of ONT assemblers must be tested using a combination of polishing strategies.

Three main methods are commonly used for scaffold ordering and orientation to generate chromosome level assemblies. Traditionally, linkage maps made of thousands of genetic markers obtained from large segregating progenies were used to anchor assembly contigs to linkage groups (Linsmith et al., 2019). However, this method can be expensive and can give false orientations due to inaccuracies in marker orientation and ordering due to genotyping errors. Synteny-based approaches can be used when a closely related high quality genome is available. However, all results obtained via these strategies are heavily biased toward the provided reference assembly, and any unique translocations or re-orderings will be lost. Further, errors in the provided reference assembly can cascade into further projects. Recently, proximity ligation methods have become a more cost effective and less biased (Peichel, Sullivan, Liachko, & White, 2017) approach for generating chromosomal level assemblies. The Hi-C method is commonly used for scaffolding genomes (Lightfoot et al., 2017; Thrimawithana et al., 2019). Hi-C data is generated by cleaving chromatin using restriction endonucleases and ligating only fragments that are close in 3D chromosomal space. The underlying premise is that the closer two fragments the more linkage markers they will share. Hi-C scaffolding algorithms take advantage of interactions at contig ends to orient and order scaffolds. However, many chromosome-level assemblies generated using Hi-C are littered with inaccurate contig placements due to shorter contigs that contain interactions spanning their entire length inhibiting Hi-C software ability to effectively orient and order these contigs accurately. Traditionally, Hi-C software are built for homozygous diploid genome assemblies and are heavily reliant on the accuracy of the reference assembly provided and many require a priori knowledge of chromosome number such as LACHESIS (Burton et al., 2013) and AllHiC (Zhang, Zhang, Zhao, Ming, & Tang, 2019). Two tools are commonly employed for Hi-C scaffolding: SALSA2 (Ghurye et al., 2019) and AllHiC (the latest version of LACHESIS). The effects of input assembly on Hi-C mapping rate and the performance of such software must also be evaluated.

*Knightia excelsa* R.Br. (rewarewa) is a nectar producing tree of the Proteaceae family, endemic to Aotearoa New Zealand. Despite its size [> 1660 species (Christenhusz & Byng, 2016)], the Proteaceae plant family has received minimal attention from genome researchers, likely due to most diversity being restricted to the southern hemisphere and the fact that the nut-producing macadamia tree is the only species in the family of significant worldwide economic interest. To date, only a partial genome assembly of *Macadamia integrifolia* has been developed, the information from which is used for genome-informed breeding (O’Connor, Hayes, & Topp, 2018). Very little genetic information is available for *K. exselsa*; however karyotype analysis indicated it is a diploid species with n = 14 chromosomes (Hair & Beuzenberg, 1958). Rewarewa is the basis of a burgeoning honey industry in Aotearoa New Zealand. Most of such honeys are produced from traditional land owned by Māori, the first nation people of Aotearoa New Zealand. Rewarewa is considered ‘*taonga*’ by Māori, meaning this tree species is treasured and under their ‘*kaitiaki*’ or guardianship, and therefore an ethical framework is necessary for managing samples and data during the project, as has been performed for other *taonga* species (Marshall et al., 2015; Morgan, Perry, & Chagne, 2019).

The objective of this research is to investigate assembly strategies using *K. excelsa* as a model (Figure 1). Subject to Māori consent, Illumina PE (61X), ONT (52X) and Hi-C data were obtained. Software for contig assembly, polishing and error correction and Hi-C scaffolding were evaluated and quality metrics measured at each step. Initial contig assemblies were generated from five long read assemblers across five subsampled sets of ONT data (reads >5 kb, >10 kb, >22 kb, >30 kb and the full data), which were corrected using a combination of long and short read polishing tools. General iterative Racon polishing followed by subsequent Pilon short read polishing was compared to a combined sequence bound and general polishing strategy of iterative Racon, Medaka and Pilon (Figure 1). The effectiveness of each ONT assembly method on chromosomal construction was assessed through Hi-C scaffolding using two software packages, SALSA2 and AllHiC, and conservation of macro-synteny against macadamia linkage maps (Alam, Neal, O’Connor, Kilian, & Topp, 2018) was tested. These tools were systematically implemented and the accuracy of each assembly was quantitatively assessed in order to identify the optimal *K. excelsa* genome assembly that could be generated from our data.

**Figure 1:**
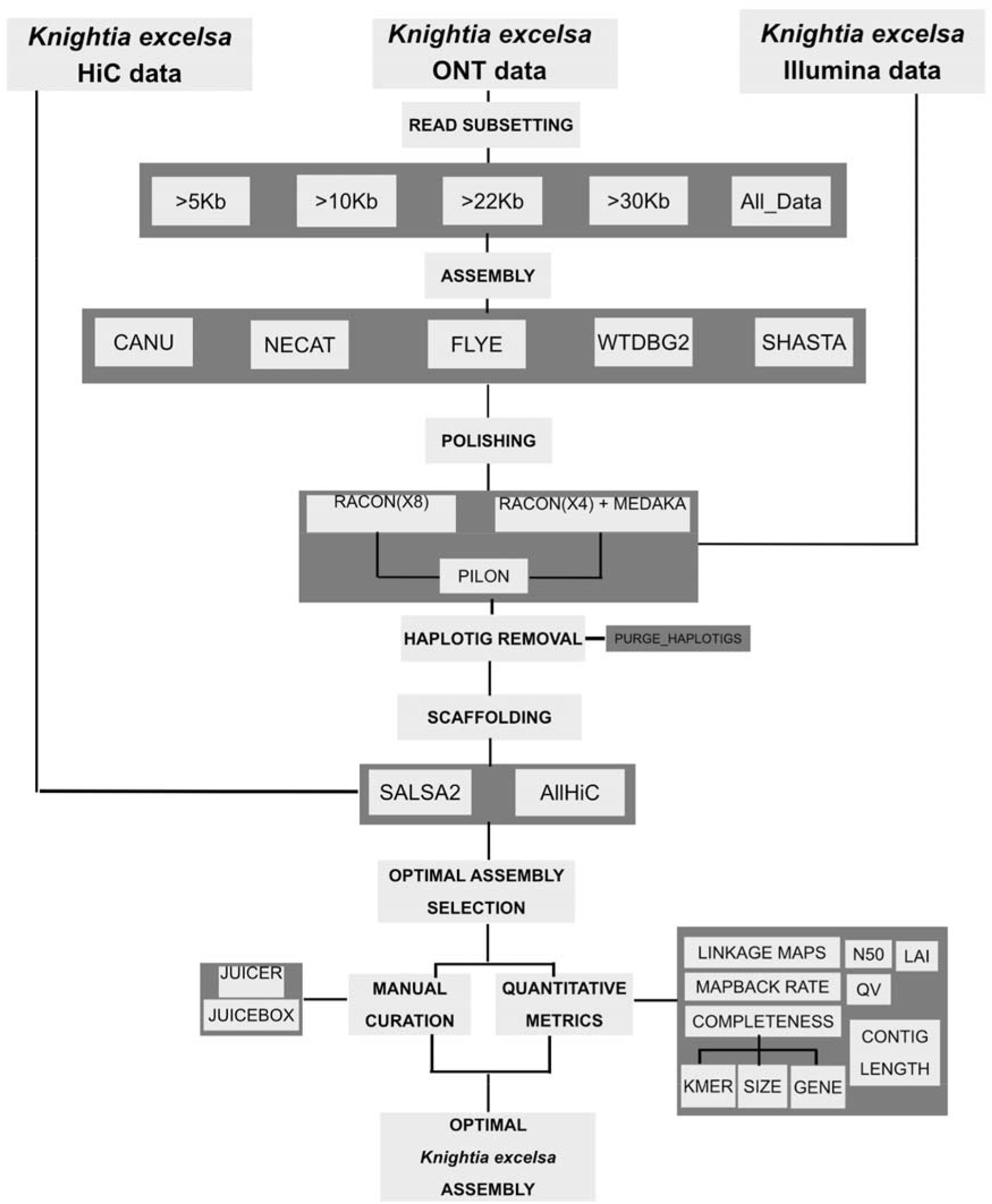
Bioinformatics workflow used for constructing an optimal *Knightia excelsa* genome assembly.

## Materials and Methods

### Sample collection

*Knightia excelsa* (rewarewa) is an endemic tree of Aotearoa New Zealand, mostly found on the North Island, and common in coastal, lowland and lower montane habitats. This evergreen tree species can grow up to 30m tall (the tallest Proteaceae), bears dark green serrated leathery leaves and dense racemes of red flowers. *K. excelsa*’s genome size was estimated to be 1.15 pg per 1C using flow cytometry.

The single *K. excelsa* tree selected for this project grows in the Warawara Forest, Northland, Aotearoa New Zealand (Lat. ± 4 m35° 22’ 1.9” South, Long. ± 4 m173° 16’ 5.4” East, altitude 440 m). The leaves were collected in November 2018, authorized by the Te Rarawa Anga Mua and the Komiti Kaitiaki for Warawara Ngahere. At the time of collection, the tree was about 3 m tall, growing in full sun, isolated from other trees, and colonizing a bulldozed site along with *Lycopodiella cernuua, Leucopogon fasciculatus*, and *Blechnum novae-zelandiae* beneath it. The tree has deep magenta flowers and was fruiting at the time of sample collection. The tree had two trunks from the same base. One trunk was 4 cm in diameter at 1.35 m above the ground and the other was 2 cm in diameter at 1.35 m above the ground. The combined cross-sectional area at breast height was 15.7 cm^2^. The leaves sampled were undamaged leaves without visible fungal infections that ranged in size from 8-12 cm long by 2-3 cm wide. Two leaf samples were collected (~20 and ~30 g).

The leaves were collected aseptically and packed in a sealable plastic bag, placed inside a Styrofoam box with crushed ice, and protected from ice burn by a stack of paper towels. The sample was delivered within two days after collection and stored at −80°C upon arrival to the laboratory.

### Nuclear genomic DNA extraction

#### Nuclei isolation

The nuclear genomic DNA was extracted from isolated nuclei as previously described (Hilario, 2018; Naim et al., 2012) with the following modifications regarding the homogenization method, the type of lysis buffer and its ratio to the amount of nuclei obtained. The leaf sample (20 or 30 g) was ground with liquid nitrogen in a precooled large mortar. The freeze/grinding cycle was repeated three times until a fine powder was obtained. The complete nuclei isolation buffer (plus sodium metabisulfite, β-mercaptoethanol and 0.5% Triton X-100) was poured in a 1 L beaker with stirrer. The powdered sample was added gradually and stirred until completely dissolved. The homogenate was filtered through two layers of Miracloth (Merck) over a funnel. The nuclei were collected by low speed centrifugation and washed twice with the nuclei isolation buffer (with sodium metabisulfite only). The final nuclei pellet was stored without any liquid at −80°C until used for DNA extraction.

#### DNA extraction

The nuclear genomic DNA was extracted with a cetyl trimethylammonium bromide (CTAB) based buffer as described(Hilario, 2018; Naim et al., 2012) with the following modifications: The isolated nuclei were lysed with 15 mL of CTAB buffer and 100 μL proteinase K (20 mg/mL). After the lysis incubation, the sample was extracted with equal volume of chloroform:iso-amyl alcohol (24:1), precipitated with ethanol and the DNA collected by centrifugation. The DNA pellet was washed with 10 mL 70% ethanol, centrifuged again and dissolved in 200 μL TE buffer. The quality of the DNA was assessed by spectrophotometry (Nanodrop) and electrophoresis separation (standard and pulse field gel electrophoresis). The amount of DNA was estimated by fluorometry (Qubit high sensitivity dsDNA kit). Average yield of nuclear genomic DNA per gram of leaf sample was 1 μg. The quality parameters were A_260/280_ = 2.0, A_260/230_ = 1.88, Qubit/Nanodrop ~ 0.5, and an average fragment size of 50 kbp.

### NGS library preparation

#### Short insert Illumina sequencing library

Eight reactions of 500 ng of nuclear genomic DNA each were set up for preparing the short insert Illumina library with the NEBNext Ultra FS II DNA library kit as described by the vendor with the following parameters: The fragmentation, end repair and deoxyadenylation incubation was 3.75 min (fragments ranging from 200 to 1000 bp). After USER digest, all the reactions were combined and split into five tubes. The library was left size selected with AMPure XP beads at 0.4X ratio followed by another left side selection at 0.2X ratio. The DNA was eluted from both bead fractions (0.4X and 0.4X/0.2X) in 30 μL TE buffer and the concentration estimated by fluorometry. A cycle test was performed with 5 ng of each size selected libraries (0.4X and 0.4X/0.2X) amplified 4, 6, 8, 10 or 12 times with NEBNext Ultra II Q5 Master mix, the Illumina universal and index primers. Ten cycles produced the optimal amplicon size after a dual size selection (0.77X/0.61X) from the 0.4X size selected library fraction (average amplicon size: 473 bp). Four reactions from this library fraction were set up under these conditions, pooled, dual size selected, quality checked and sent to our service provider (Custom Science, New Zealand) to be sequenced.

#### Long range sequencing library (Hi-C)

The long range Hi-C sequencing library was prepared with isolated nuclei as starting material. The nuclei enrichment method is similar to the protocol described above but with extra steps to remove contaminants and large particle debris with polyvinylpolypyrrolidone (PVPP) and Percoll^®^ gradients, respectively. The Hi-C library was prepared with a combination of kits and in-house methods. The nuclei crosslinking, quenching, washing, lysis and chromatin normalization steps were performed according to the Dovetail Genomics Hi-C kit. The chromatin lysate was bound to AMPure XP beads and washed with 5 sets of 1 mL Wash buffer (Dovetail Genomics Hi-C kit). Chromatin fragmentation and biotinylation were performed with the Fragmentation buffer and Fragmentation Enzyme mix from the Phase Genomics Hi-C kit for plants Version 1.0. Once the digestion was completed, the captured chromatin was washed twice with Wash buffer (Dovetail Genomics). The intra-molecular ligation was performed in 500 μL of 1X T4 DNA ligase buffer (Invitrogen) and 10 units of T4 DNA ligase (Invitrogen). The ligation was performed at 16°C in a thermomixer (Eppendorf) at 1250 rpm overnight. The ligation mixture was discarded and the crosslink reversal was performed by adding 50 μL 1X CutSmart buffer (New England Biolabs) and 20 μg Proteinase K (Qiagen) and incubated at 55°C for 15 min followed by 45 min at 68°C at 1250 rpm. The released DNA was transferred to a new tube and purified with AMPure XP beads at 2X ratio. The DNA was eluted in 150 μL 10 mM Tris-HCl pH 8 and the biotinylated molecules captured with Dynabeads M280 (Invitrogen) according to the manufacturer’s protocol but using150 μL Bead Binding buffer (Phase Genomics) for coupling the biotinylated molecules to the beads and continue with the Phase Genomics Hi-C kit for plants protocol. The amplified library was size selected by agarose gel electrophoresis followed by an AMPure XP double size selection (0.77X/0.64X). The average fragment size of the selected amplicons was 500 bp. The size selected amplicons were assessed by capillary electrophoresis (Fragment Analyzer) and showed an average fragment size of 441 bp, at 1.5 ng/μL and 4.7 nM. The amplicons were sequenced (150 b paired end reads) and delivered 221,731,503 raw PE reads, and 66.96 Gb.

#### PromethION Oxford Nanopore Sequencing

The PromethION libraries were prepared by the contracted service provider (Custom Sciences, New Zealand) with ~ 50 μg of nuclear genomic DNA preparation described above.

##### Genome assembly and assessment

###### Initial quality assessment and subset generation of Oxford Nanopore reads

All datasets were base-called using Guppy flip flop software package (**Supplementary Material 1**) and quality assessed using the FASTQC raw reads for quality assessment. ONT sub-setting was carried out using the porechop software package. The data was subsampled by read length into five read sets: >5 kb reads only (52x), >10 kb reads only (50x), >22 kb reads only (33x), and > 30 kb (23x) reads only, and all data. These values were selected in order to retain sufficient sequencing depth within each subset.

###### Oxford Nanopore Assembly

Five long read assemblers were used: CANU, FLYE, WTDBG2, SHASTA and NECAT (for parameters and versions used see **Supplementary Material 1**). In order to further understand the effects of polishing strategies on assembly accuracy combinations of polishings methods were examined and haplotigs purged [41]. These include general polishing strategies: Racon with four rounds (RX4) of polishing only, Racon with eight rounds of polishing both before (RX8) and after pilon (RX8_SR) polishing and haplotig purging (RX8_SR_PH). A Sequencer specific strategy alone was also included: Medaka only (M) polishing, as well as combined polishing approaches: Medaka with four iterations of Racon polishing both with (M_RX4) and without pilon polishing (M_RX4_SR) and haplotig purging (M_RX4_SR_PH). Each assembly was initially quality checked using QUAST, BUSCO and LAI.

###### Hi-C mapping

The Hi-C dataset was filtered using the Phase Genomics filtration guidelines (link). The data successfully passed all quality assessment analysis requiring no additional filtration. The data was mapped to each generated ONT contig set using bwa mem and scaffolding was carried out by SALSA2 and AllHiC (see **Supplementary Material 1** for parameters and versions used).

###### Hi-C assembly quantitative quality assessment

Each Hi-C assembly kmer spectra profile was assessed through meryl and consensus accuracy and completeness analyzed using the merqury toolkit. Map back rates were also used to assess the quality of each assembly using samtools (Cock, Bonfield, Chevreux, & Li, 2015) flagstat statistics (see **Supplementary Material 1** for parameters used). All assemblies were additionally compared using the LAI index which assesses the LTR repeat completeness of plant genomes specifically. On top of this, assemblies were compared through QUAST (Gurevich, Saveliev, Vyahhi, & Tesler, 2013) and BUSCO. Along with contact map manual inspection by PretextMap and PretextView (https://github.com/wtsi-hpag/PretextView).

###### Utilization of *Macadamia* linkage maps for QC

Nine linkage maps accompanying 64bp DartSeq reads were downloaded from the Southern Cross University data repository [http://dx.doi.org/10.25918/5dc2589924ca2]. These reads were aligned using blastn to three Hi-C assemblies (the best assemblies selected based on quantitative assembly accuracy metrics) and only unique hits of > 90% identity were used. These markers were mapped to each assembly using ALLMAPs (H. Tang et al., 2015) (see **Supplementary Material 1** for versions and parameters).

###### Computational Resources

The majority of analyses were carried out on the New Zealand eScience Infrastructure high performance computer on the Mahuika partition. The Mahuika partition consists of a Cray CS400 Cluster High Performance Computer with 8,424 x 2.1 GHz Intel Broadwell cores and 30 Terabytes of memory along with IBM ESS Disk and SSD storage. For computational efficiency each assembly was run using minimal requirements (See **Supplementary Material 1**). CANU Assemblies were performed at the University of Otago’s Biochemistry Servers, which have 1 Tb of memory, and 8x Intel(R) Xeon(R) CPU E7-8860 v4 with 18 cores each, and 2 threads per core.

## Results

### Sequencing data

In total, 2.3M ONT sequencing reads were obtained totaling 52.5 Gbp of data and with a read N50 of 28kbp. Table 1 indicates the initial summary read statistics for the ONT data. The significance of basecalling was assessed both before and after base-calling using MinIon QC (Lanfear, Schalamun, Kainer, Wang, & Schwessinger, 2019) (**Supplementary Material 2**). A significant increase overall Q score was achieved and specifically for longer read lengths. Hi-C data produced from the Phase Genomics kit and Illumina PE sequencing yielded 443M reads in total (67Gb of data). Short read WGS data was obtained and consisted of 407M PE Illumina reads. Through kmer counting (k=21) a genome size of 0.95Gb and heterozygosity as 0.1-1.0% was estimated.

**Table 1:** Basic Statistics of Oxford Nanopore Technologies sequencing data for *K. excelsa*.

### An in depth performance critique of ONT assemblers and iterative polishing

An assessment of the performance of five long read assemblers; NECAT, CANU, SHASTA, FLYE and WTDBG2 was carried out and initial contig sets generated. ONT data was then split into five subsamples based on read length, these subsamples included >5 kb, >10 kb, >22 kb, >30 kb and all read lengths. Assembly performance was compared across these subsamples to assess how read length might affect the performance of individual long read assemblers.

Firstly, the output from iterative long read polishing using Racon was examined to explore potential effects on assembly accuracy. Based on contiguity, total length and N50, eight rounds of iterative Racon polishing had little effect on the accuracy of the initial contigs sets across CANU and FLYE with quantitative metrics remaining consistent regardless of number of polishing iterations applied across each read subsample (Figure 2). Interestingly, the NECAT assembly generated for All_Data subsample appears collapsed after two rounds of polishing, with a drastic reduction in contig number and total assembly length below flow cytometry estimations (1 Gbp). NECAT failed to complete for all other read length subsamples and therefore was not included in further performance comparison analysis. Shasta-generated assembly metrics remain consistent in the >10, >22 and >30kb read subsamples with iterative polishing having little effect, however, total genome length slightly reduced in >5kb and All_Data subsamples. WTDBG2 assembly metrics remained robust against polishing for all read subsamples, apart from the >10kb subsample which encountered a total length expansion.

**Figure 2:**
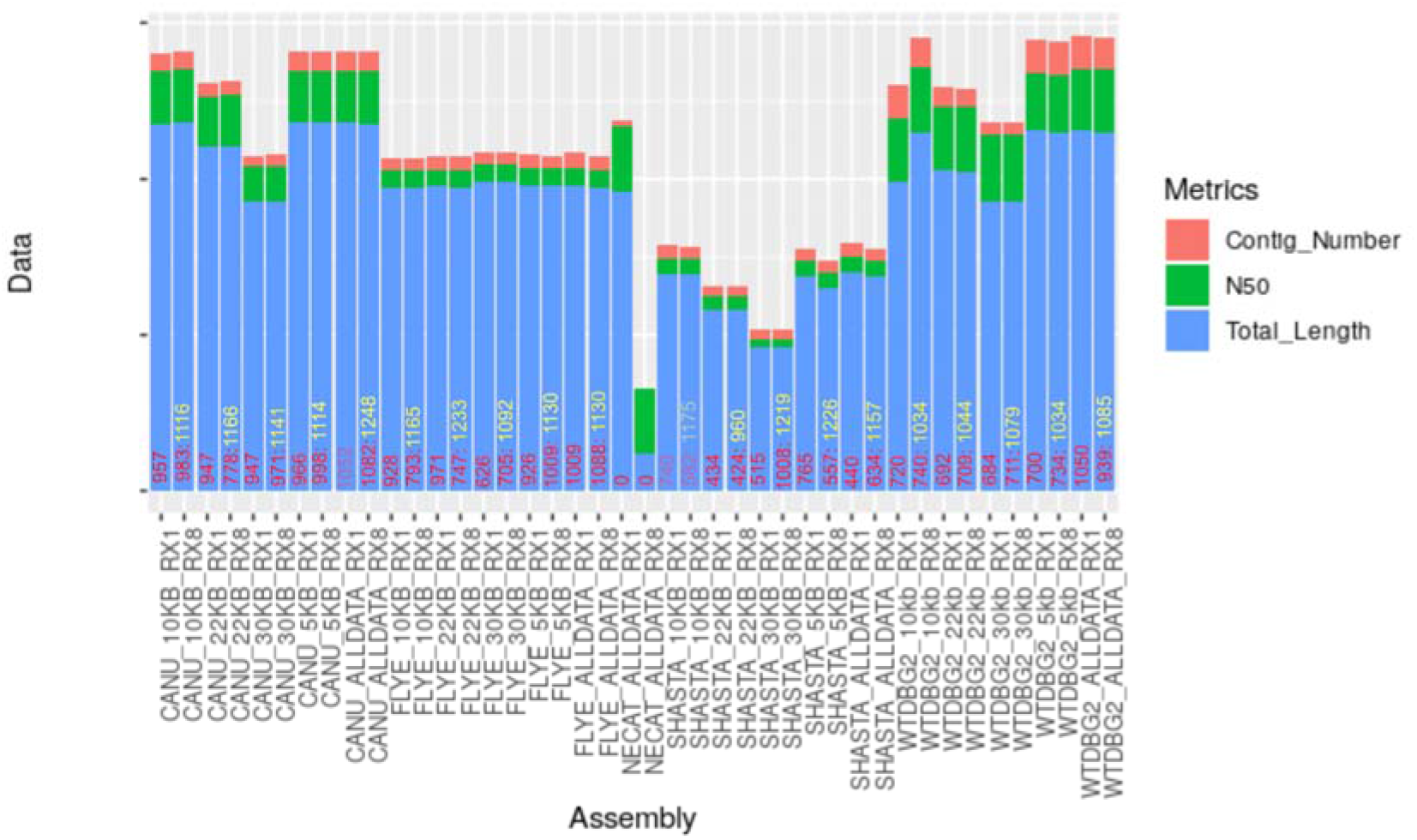
A comparison of the performance of five ONT assemblers and iterative polishing by Racon across assemblies generated by all read length subsamples. A comparison of contig number, N50 and Total length of a single Racon polishing in comparison to eight rounds of polishing. Complete gene scores after long read polishing are displayed in red inside its corresponding assembly bar and numbers in yellow indicate complete gene scores post short read Pilon polishing.

Gene completeness was assessed; generally, iterative long read polishing slightly increased the number of complete genes identified across assemblers, except for CANU and FLYE’s >22 kb subsample assembly that experienced a 169 and 224 complete gene reduction, respectively, and FLYEs >10 kb assembly that showed a 135 reduction. SHASTA experienced a complete gene reduction of 158, 10 and 208 in >10 kb, >22 kb and >5kb read subsamples, respectively, and WTDBG2 only experienced a reduction of 111 genes in the assembly constructed by the All_Data subsample. Across all assemblies the accuracy of Illumina data increased the gene completeness through Pilon polishing. Interestingly, the ONT assemblies that experienced a reduction in gene completeness score after iterative long read polishing incurred the greatest increase in score post short read polishing, with CANU and FLYE >22 kb subsample experiencing an increase of 388 and 286 genes, respectively, and SHASTA >5 kb, >10 kb and >22 kb read subsamples gaining 669, 593 and 536 genes, respectively.

In terms of total length and contiguity, FLYE’s performance appeared the most robust, with total length and N50 values remaining consistent (Figure 2), however, contiguity was increased in the >30 kb subsample (smaller number of contigs). WTDBG2, SHASTA and CANU appeared to perform much better with longer read lengths, based on the lower number of contigs and increased N50 for the >30 kb subsample. However, the total length of the assembly seemed to be compromised, with a decreased total length below the expected genome size when using the >30 kb subsample. This may have been the result of low depth of coverage in this read length subsample and may be improved with an increase in data volume within this read length subsample.

### An assessment of post-assembly polishing strategies across read subsets

In the previous section it was clear that iterative Racon polishing improved assembly metrics. The assemblies were also used to assess polishing strategies (Figure 3). The first polishing strategy implemented eight iterative rounds of long read polishing by Racon (RX8), with the results after four rounds are displayed (RX4) in Figure 3 for comparative purposes, followed by an additional short read polishing by Pilon (RX8_SR) - an example of a “General” polishing strategy. The haplotigs from each contig set were then removed using the *purge_haplotigs* software package (RX8_SR_PH). This experiment aimed to investigate the benefits of using Racon, a tool developed for assemblies prior to consensus, on assemblies after consensus has taken place. The second polishing strategy designed consisted of Medaka (M) only polishing in order to test a tool developed specifically for consensus polishing and an example of a “Sequencer specific” polishing strategy. The third strategy integrated both tools through an initial four rounds polishing with Racon, followed by a single round of both Medaka (M_RX4) and Pilon polishing (M_RX4_SR) that was then purged of haplotigs (M_RX4_SR_PH). The effect of these polishing strategies was assessed by examining the total read assembly size, contiguity and BUSCO completeness (Figure 3).

**Figure 3:**
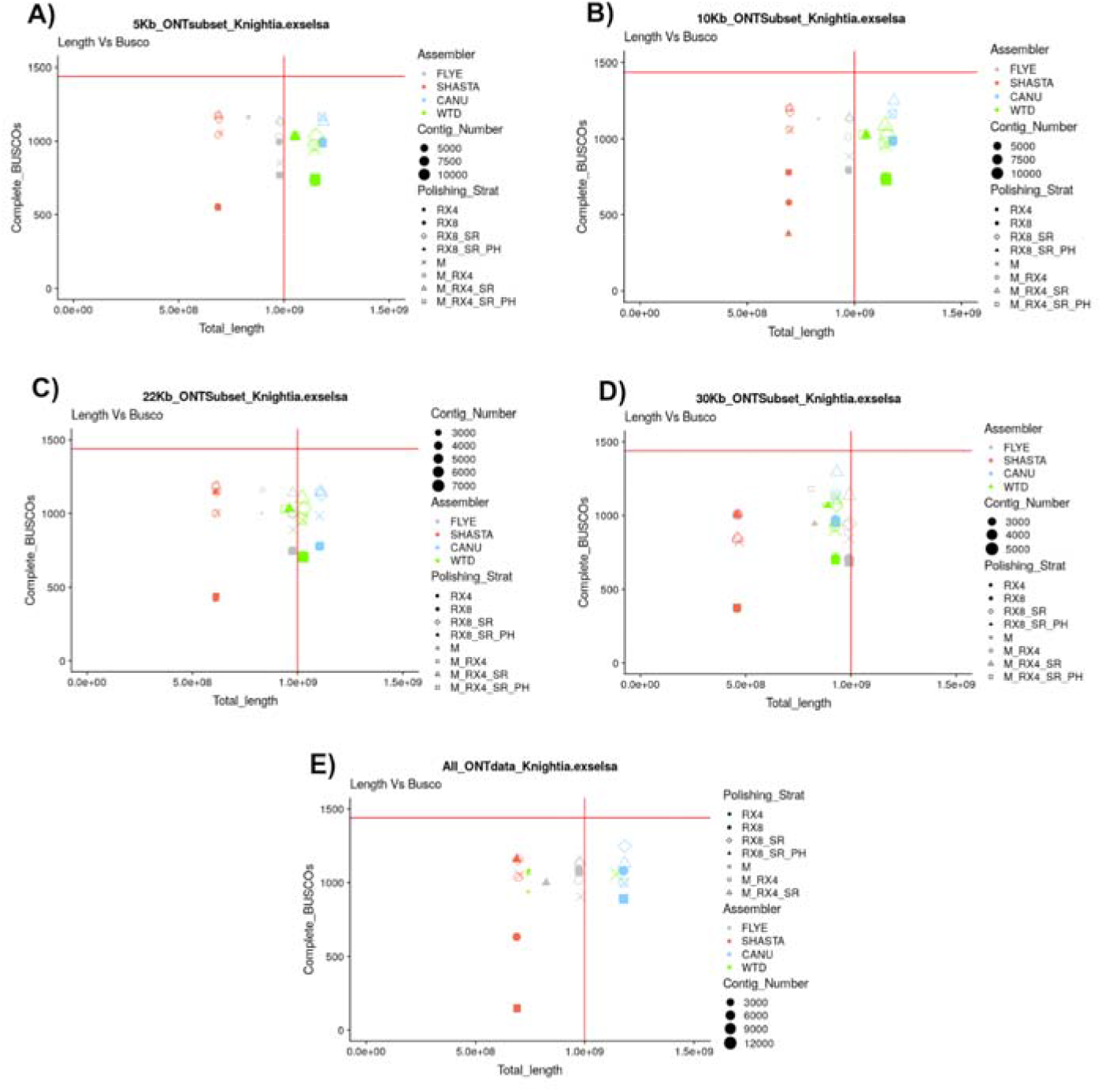
A comparison of polishing strategy performance on all read subsets using five read subsets across contig sets generated by four long assemblers. A) >5kb read length subsample, B) >10kb read length subsample, C) 22kb read length subsample, D) >30kb read length subsample, and E) All_Data read length subsample. Vertical red line illustrates the estimated genome size, horizontal red line highlights maximum number of BUSCO genes within the *embryophyta_odb9* dataset. Increasing data point size indicates an increase in contig number within the assembly.

With the >5 kb and >10 kb subsamples, CANU, FLYE, WTDBG2 and SHASTA perform consistently well in regards to total length in strategies combining Racon and Medaka (M_RX4, M_RX4_SR_PH and M_RX4_SR) (Figure 3a,b). Here, both FLYE and WTDBG2 obtained a contig set that was the most representative of the expected genome size, with no bias toward general or sequencer specific polishing identified. However, when gene completeness is considered all assemblers showed a benefit from using polishing strategies that incorporate Medaka (M, M_RX4, M_RX4_SR_PH and M_RX4_SR).

Analyses using the >22 kb subsample indicated that read length and depth of coverage enabled the WTDBG2 assembler to more accurately represent the total genome size while retaining gene completeness in comparison to that with the >5 kb and >10 kb subsamples and again no polisher bias was apparent (Figure 3c). However, they remained more fragmented when compared to those constructed by FLYE and CANU, whose bias toward a combined polishing (M_RX4, M_RX4_SR_PH and M_RX4_SR) remained consistent to that constructed with >5 kb and >10 kb subsamples.

CANU outperformed all other assemblers in regards to gene completeness when only read lengths of >30 kb were included and compared to its performance in all other read length subsamples, and there is a clear bias towards polishing strategies that include Medaka. Here, both SHASTA and WTDBG2’s performance were compromised from the lack of read depth, as highlighted by the continual reduction of total assembly size as read length cut offs increase and both assemblers remain consistent across polishers. FLYE’s performance at this read length remained consistently biased toward Medaka incorporated strategies (Figure 3d).

When all data is included, SHASTA and WTDBG2 failed to represent an accurate total length despite becoming more contiguous. Both assemblers experienced the same shortcomings when polished using Racon only, Medaka only and combined strategies. CANU’s total genome length suffered despite good performance in regards to gene completion and retained its bias towards Medaka polishing. FLYE’s performance was relatively constant across all subsets. Interestingly, although an integrated Racon/Medaka polishing strategy still performed better here, Racon only polished assemblies performed better than across other read length subsamples (Figure 3e).

Assemblers performed differently to haplotig purging, with diploid aware assemblers such as CANU experiencing a reduction in gene completeness score when this technique was implemented across all read subsamples. Overall, FLYE retained gene completeness and increased its contiguity in response to haplotig purging, however, across each read subset the total genome size experienced a reduction to below what was expected for the genome. WTDBG2 responded similarly to FLYE across all read subsets whilst SHASTA experienced varying shifts in performance over all read subsets.

Across both “general” and “sequencer-specific” polishing, WTDBG2 assemblies appeared the most fragmented, with expanded genome total lengths and no bias toward polishing strategy was identified. In contrast all SHASTA assemblies appeared highly contiguous, however, they had unexpectedly small total lengths and are unaffected by polisher. FLYE and CANU appeared to perform best with FLYE performing equally well across all polishing methods in regards to genome size representation. These results highlight a clear bias toward polishing strategies that incorporate Medaka as opposed to those utilizing Racon alone, this indeed suggests that polishing methods specific to the sequence platform utilize have a superior performance than “General” polishers that are not platform specific.

### Comparison of Hi-C scaffolding strategies in preparation for pseudo-chromosome construction

Eighteen initial ONT assemblies were generated utilizing the All_Data read length subset, however, it was not known which subsample was the most appropriate for Hi-C mapping or what Hi-C mapping workflow to use. Hi-C data was mapped and duplicates filtered in accordance with Phase Genomics quality assessment guidelines (**Supplementary Material 3**). Each assembly assessment indicated that the library preparation would sufficiently inform the underlying assembly. However, for *de-novo* assembly scaffolding high quality read pairs between contigs are crucial and those found within contigs are uninformative. During QC it was found that WTDBG2 and NECAT assemblies had a reduced intercontig RP percentage when compared to that of CANU, FLYE and SHASTA each peaking at 19%, 21% and 22%, respectively.

All ONT assembly constructs created were scaffolded using two software packages, SALSA2 and AllHiC. After initial scaffolding a series of quantitative quality assessments including gene completeness, total length assessment, map back rate, kmer spectra analysis, kmer completeness profiling, consensus accuracy and LAI were calculated.

Through comparing quantitative metrics (Figure 4), the quality of assemblies with a poor initial contig set, specifically in regards to total length accuracy such as initial NECAT contigs, could not be recovered by either Hi-C scaffolding pipelines. This holds true in WTDBG2 contigs sets, with total genome size and overall kmer completeness scores falling short after Hi-C data was mapped. CANU assemblies performed well, furthermore scaffolding on assemblies based on a Racon/Medaka/Pilon polishing strategy outperformed those produced by Racon/Pilon only polishing. CANU initial ONT assemblies produced scaffold sets by both AllHiC and SALSA2 that were 93% kmer complete. However, read map back rates suggested SALSA2 utilized 7% more input data when compared to scaffold sets produced by AllHiC. The CANU-SALSA2 strategy also outperformed in regards genome completeness with scaffold sets containing 77% complete genes in comparison to CANU-AllHiC scaffolds having only 70% complete genes. All FLYE scaffold sets perform well in regards to kmer completeness. However, FLYE-SALSA2 and FLYE-AllHiC mappings with Racon/Pilon only polishing produced more accurate total genome lengths but had lower gene completeness scores. By comparing metrics across the eighteen initial assemblies, two assemblies were selected for further analyses before pseudo-chromosome construction - those produced by the FLYE/Medaka/Racon/Pilon/SALSA and FLYE/Medaka/Racon/Pilon/AllHiC workflows.

**Figure 4:**
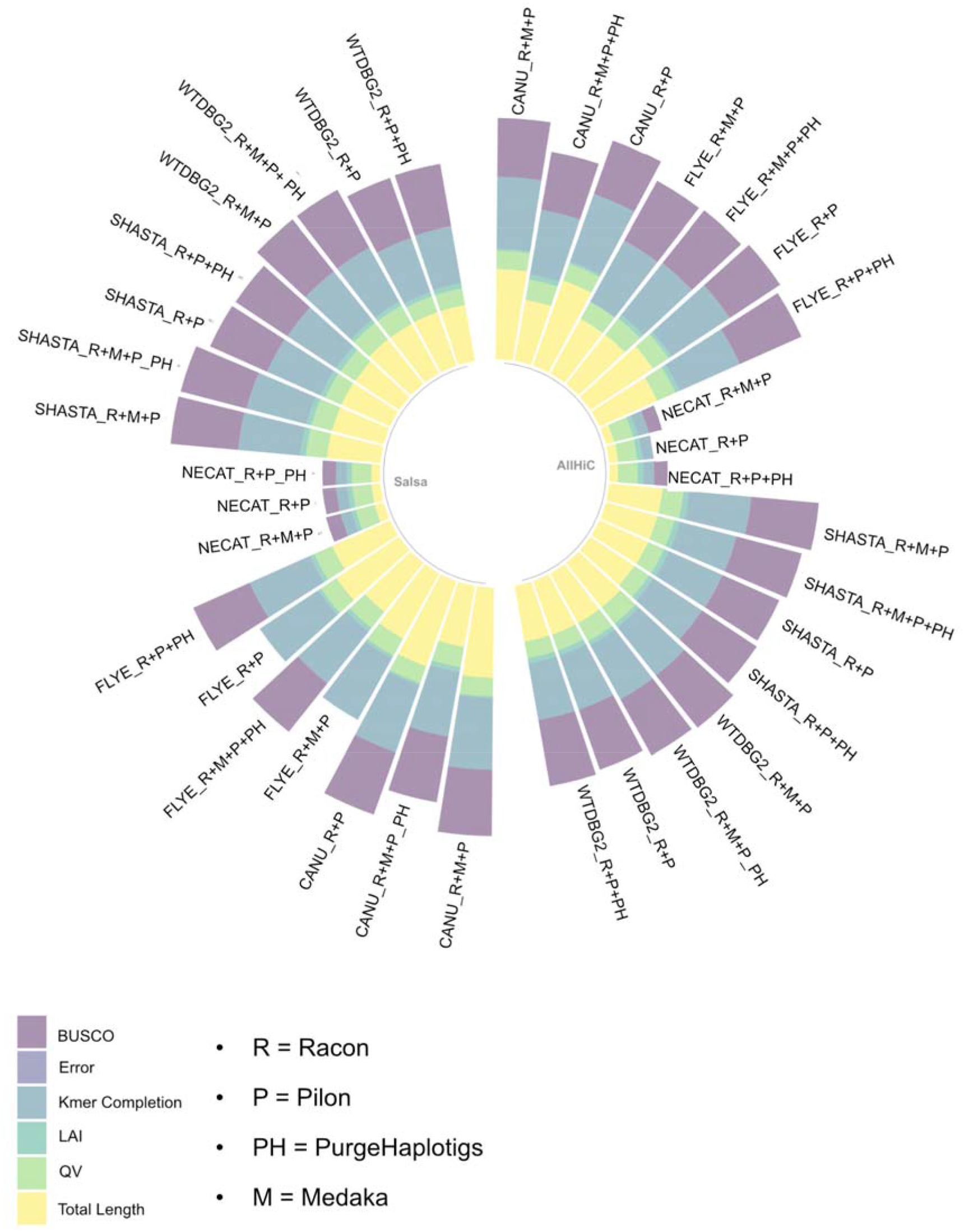
An extensive quantitative quality assessment of scaffold sets produced by both SALSA2 and AllHiC on ONT assemblies constructed from all read lengths. Quantitative metric summary of Hi-C assemblies generated using AllHiC and SALSA 2. Figure displays LAI, kmer completeness, base error rate, consensus accuracy (QV), total length, and gene completeness (BUSCO).

### A comparison of Hi-C scaffolding strategies on ONT assemblies generated by read length subsampling

To assess the impact of the underlying read length, coverage and genome assembly quality on Hi-C scaffolding each subsampled (>5 kb read, >10 kb read, >22 kb read and >30 kb read samples) ONT assembly (Racon+Pilon polished assemblies only) was taken and further scaffolded by AllHiC and SALSA2. Each scaffolded assembly was quality assessed as summarized in Table 2.

**Table 2:** Quantitative quality assessment of the impact of two Hi-C mapping strategies on five read length subsampled genome assemblies of *Knightia excelsa* produced across four long read assemblers.

**Table 3:** Pseudo-chromosome length of the final *K. excelsa* genome assembly.

The results (Table 2) demonstrated the inability of Hi-C scaffolding to effectively reduce the high contig number found across all WTDBG2 initial ONT assemblies. AllHiC failed to complete scaffolding on the >5 kb and >10 kb read length subsamples and only a 16% and 17% reduction in contig number was achieved for >22 kb and >30 kb subsamples, respectively. AllHiC-WTDBG2 >30 kb and >22 kb subsample assemblies consisted of a single ‘mega’ scaffold that contained >97% of the total length (Figure 5d), which is not consistent with the expected karyotype for *K. excelsa*. This peculiar length distribution was rectified in the All_Data read subsample (Figure 5a). Despite producing less contiguous assemblies, SALSA2 scaffolding produced using the initial WTDBG2 assemblies appeared more accurate with optimal performance resulting from using the >30 kb subsample, achieving a kmer completeness value of 83%, gene completeness score of 85% and an 85% reduction in contig number observed, and gave a contig length distribution (Figure 5c) more similar to the known karyotype.

**Figure 5:**
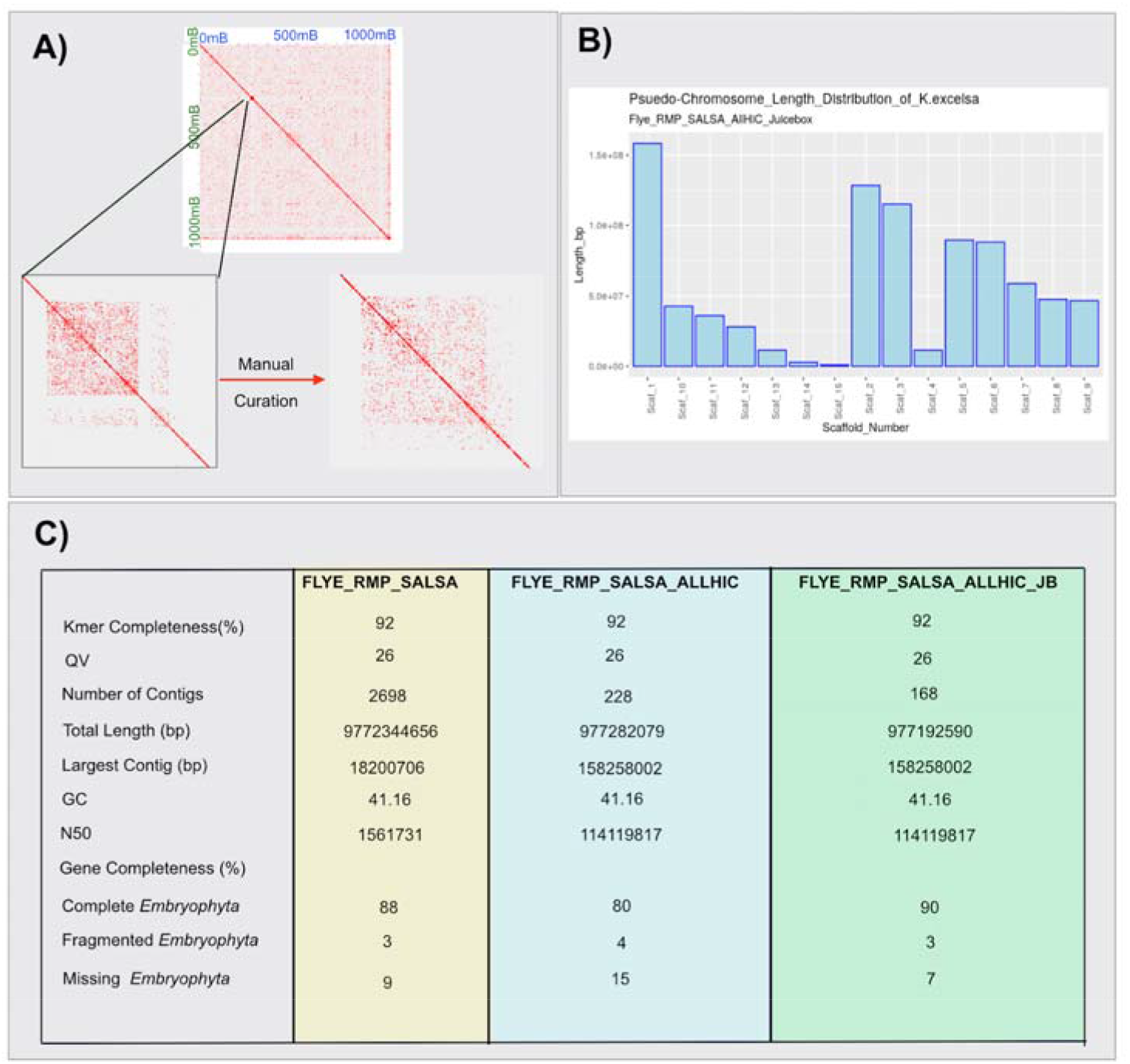
A comparison of scaffold lengths for *K. excelsa* genome assemblies produced by both SALSA2 and AllHiC Hi-C mapping. A) Scaffold lengths for assemblies generated using read length subsampled data by four long read assemblers across four read lengths and Hi-C data mapped using SALSA2. B) Scaffold lengths for genomes produced using data subsampled by read length by four long read assemblers and Hi-C data mapped using AllHiC. C) Scaffold lengths for assemblies produced using all read length data by five long read assemblers utilizing two alternative polishing strategies and Hi-C data mapped using SALSA2. D) Scaffold lengths for assemblies produced using all read lengths by five long read assemblers utilizing two alternative polishing strategies and Hi-C data mapped using AllHiC.

Similarly, the problem of greatly reduced SHASTA ONT assembly length relative to the genome size estimated by flow cytometry, was not resolved by Hi-C scaffolding using either SALSA2 or AllHiC. AllHiC scaffolding, although greatly reducing contig number produced a suspicious ‘mega’ scaffold similar to the distribution found in WTDBG2 scaffolded assemblies (Figure 5d). However, unlike WTDBG2 scaffolds the issue for SHASTA scaffolds was not resolved in the All_Data read subsample. Again, SALSA2 scaffolds constructed using the >30 kb subsample appeared to be the most accurate, with a gene completeness of 85% and a 99% reduction in contig number and no ‘mega’ scaffold (Figure 5c). Unfortunately, due to the poor total length of the initial ONT assembly provided by SHASTA, the kmer completeness score of the final assembly remained low at 55%, therefore drastically under-utilizing the data provided.

AllHiC performed optimally when utilizing the more robust initial ONT assemblies generated by FLYE (Table 2), achieving a >90% contig number reduction across all read length subsamples, and although a ‘mega’ scaffold still persists, its size reduced to ~50% of the total genome length and the distribution seemed comparable to SALSA2 scaffolds using All_Data subsample (Figure 5a and Figure 5b). Interestingly, the >5 kb and >10 kb subsampled assemblies represented more of the data, with higher gene completeness values and a kmer completeness score of 91% for both, in comparison to >22 kb and >30 kb subsamples that had lower gene completeness scores and a kmer completeness value of only ~80%. Comparatively, SALSA2 scaffolding failed to reduce contig numbers in the >5 kb and >30 kb subsamples and only achieved a 34% and 36% reduction in >10 kb and >22 kb read subsamples, respectively. Although SALSA2 assemblies have lower contiguity in comparison to that of AllHiC, they consistently outperformed in terms of gene completeness.

Finally, AllHiC scaffolding for all CANU ONT assemblies yield a suspicious contig length distribution, with >92% of total genome size being placed on a single mega-scaffold. Gene completeness values were also poor with all samples below 80%, which was surprising considering the scaffolds generated were the most kmer complete, peaking at 93% across samples. Again, the optimal initial CANU assembly from the >30kb read subsample generated the optimal assembly after SALSA2 scaffolding, with gene completeness of 85% and 93% kmer completeness. In this case, SALSA2 also reduced contig numbers by 39%.

Overall, AllHiC with prior knowledge of karyotype performed well with higher contiguity and longer scaffolds. However, on comparison of the scaffold length distribution, SALSA2 generated more uniform scaffold lengths across subsamples whereas ALLHiC tended to generate assemblies with one single mega-scaffold and a multitude of much shorter scaffolds - which is not in agreement with the known karyotype of *K. excelsa*.

### Verification of Hi-C scaffolding using synteny with nine Macadamia genetic maps

After quantitative metric assessment along with scaffold length distribution analyses of assemblies across all readlength subsamples, two All_Data assemblies generated using the initial ONT workflow of FLYE/Racon/Medaka/Pilon and further scaffolded with SALSA2 or AllHiC were selected for further validation using 14 linkage groups generated for the macadamia genome (Nock et al., 2020). Macadamia was selected as it belongs to the Proteaceae and shares a karyotype of 14 chromosomes with *K. excelsa*. Using BLASTn, unique markers were identified (mean = 227 unique markers identified per map) across nine maps. These unique markers were then mapped using ALLMAPs (H. B. Tang et al., 2015) to the two *K. exselsa* assemblies and the order and orientation of scaffolds were visually examined for synteny (**Supplementary Material 4**). Whole genome alignments were constructed to compare each Macadamia informed assembly to its original Hi-C assembly. From this it was clear that scaffolds generated using SALSA2 shared a greater proportion of synteny with Macadamia when compared to AllHiC scaffolds, although they were less contiguous. To further assess accuracy, the location of the telomere motif “TTAAGGG” was identified in each assembly using EMBOSS (Rice, Longden, & Bleasby, 2000) and visual constructions created using ChromoMap (Anand, 2019) (**Supplementary Material 4**). These analyses indicate that both Hi-C scaffolders are unable to accurately represent telomere sequences, however, SALSA2 scaffolds generated a more accurate assignment than scaffolds constructed using AllHiC.

Overall the analyses confirmed that SALSA2’s scaffolding of the initial FLYE ONT assembly and polished with a combined Medaka/Racon/Pilon strategy outperformed the scaffolds produced by AllHiC, using the same initial ONT assembly, with regard to the orientation and ordering of scaffolds and accuracy of regions of complexity.

### Pseudo-chromosome level assembly construction

After linkage group validation the SALSA2-FLYE based assembly was selected for further scaffolding using the AllHiC package. Here, pseudo-chromosomes were identified and metrics of 92% kmer completion, a QV score of 26 and a total length of 0.97Gb were obtained. However, although a higher level of contiguity was achieved, gene completeness scores dropped to 80% from 87%. On inspection of the AllHiC contact map a mis-assembly was identified and rectified through manual intervention using JuiceBox (Figure 6a). This manual curation resulted in a 10% increase of gene completeness whilst retaining the contiguity provided by AllHiC. The final *K. excelsa* assembly had a 90% and 97% gene complete using the *embryophyta & eukaryota* databases, respectively, an N50 of 114Mb (Figure 6c), a karyotype similar to that expected for the species (Figure 6b) and is available through https://doi.org/10.7931/paqg-kk20.

**Figure 6:**
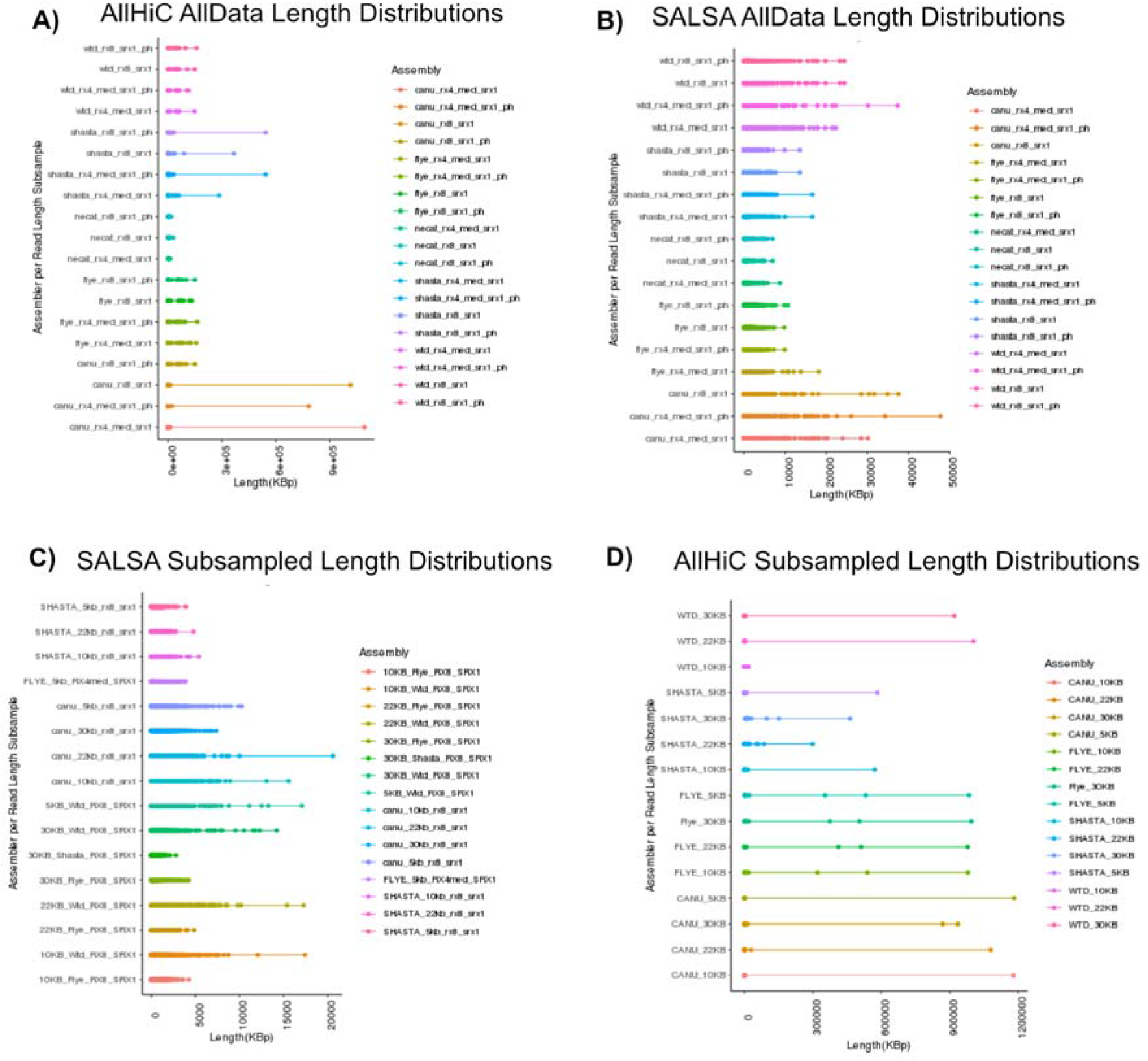
Pseudo-chromosome assembly curation and validation using both quantitative metrics, karyotype evaluation and manual curation. A) Illustrates the contact map generated for FLYE/Racon/Medaka/Pilon/SALSA2/AllHiC assembly and zooms in on the mis-assembly both before and after manual correction. B) Illustrates the scaffold lengths of the 13 pseudo-chromosomes and the longest two additional scaffolds. C) A panel of quantitative statistics generated to compare each scaffolding iteration.

## Discussion

### Long Read Assembler Performance with Iterative Polishing

Here we report and outline the importance of utilizing a synergistic assembly, polishing and Hi-C scaffolding workflow, using the construction of the ‘optimal’ pseudo-chromosomal assembly for *K. excelsa* as the exemplar.

It is crucial that an appropriate assembler and sequencing strategy is selected prior to data generation, in order to maximize the use of both the data type and data volume during assembly to meet the goal of constructing an assembly optimal for individual project needs. The performance of five ONT assemblers across five read length subsets [reads >5 kb only, >10 kb only, >22 kb only, >30 kb only and the entire data (unfiltered by length)] was investigated. This work was used to construct an optimal ONT initial assembly for further Hi-C scaffolding but also has the potential to inform future project’s assembly workflows where data volume is different, saving time and computational resources spent on benchmarking. In order to compare performance across assemblers quantitative metrics such as total length, contiguity and N50 for each assembly were generated.

SHASTA was built for quick assembly constructions and was originally developed for the human genome, with 11 human genomes assembled in nine days on a single compute node. This feat was made possible by strategic read length encoding, reduced marker representation and heuristics. However, although being fast, SHASTA requires a large amount of RAM, 1-2 Tb for the human genome (K. Shafin et al.), which is not always readily available and compromises assembly accuracy over contiguity and total length. Figure 2 highlights this assembler shortcoming with assemblies across all read lengths unable to represent the expected total length of *K. excelsa* (1Gb) whilst consistently achieving high gene completeness scores.

Interestingly, Figure 2 illustrates this assembler’s dependency on a high depth of coverage for optimal performance, with subsamples that include additional read depth, e.g. >5 kb assemblies and All_Data being of better quality than those with a lower depth of coverage such as >30 kb. SHASTA appears to positively respond to iterative Racon polishing with continuous improvements in contiguity found across subsamples when implemented (Figure 2). When alternative polishing strategies were tested across subsampled datasets SHASTA appeared unbiased with both “general” and “sequencer-specific” polishers performing equally well in the All_Data, >30Kb and 22Kb subsamples whilst Medaka based approaches seem to outperform “general” approaches in the >5 and >10Kb subsample.

WTDBG2 produces highly fragmented assemblies in comparison to all other assemblers when shorter read lengths are included, with enhanced N50 and reduced number of contigs occurring when only longer read lengths are provided (Figure 2). This result is due to its underlying algorithm having only a single consensus step and is reiterated by a significant improvement of contiguity, gene completeness and N50 after Racon and Pilon polishing, which has been identified as an issue in other plant species assemblies such as *Acer yangbiense* (Yang et al., 2019).

Consistent with results found for prokaryotes (Wick & Holt, 2019), we demonstrated WTDBG2’s decreased performance at lower read depths with assemblies produced by >30 kb read subsample failing to span the expected total length, which was not rescued by iterative polishing. WTDGB2 was the only assembler identified without a bias toward a Medaka-based polishing strategy, with Racon-based strategies also performing well.

CANU’s optimal performance is reached when only longer read lengths are provided, and performance is compromised when additional shorter read length data is added. This clearly shows a preference by this assembler for read length over depth of coverage, supporting claims made by the developers that only >20X coverage is required for accurate assembly. Algorithmically, CANU contains extensive rounds of error correction and consensus, and the developers do not suggest additional long read polishing. Thus, as expected, the post assembly iterative polishing shown in Figure 2 has the least effect on these assemblies when compared to all other assemblers, as the initial assemblies generated have substantially fewer errors to correct. This finding does not appear to be specific to Racon and Medaka long read polishers only, as these minimal effects have also been identified by other long read polishing tools. For example, it has been shown that by polishing bacterial assemblies generated by CANU using Nanopolish an increase in errors found in the assembly occurred when compared to short read polishing alone (Goldstein, Beka, Graf, & Klassen, 2019). Similar results are represented in Figure 2, as implementing the short read polishing recommended by developers achieved a substantial gene completeness score across all subsamples.

FLYE achieved the most robust performance, with the assemblies generated not significantly impeded by the addition or exclusion of certain read lengths or read depths. This result has been demonstrated for bacterial genome assemblies whereby the assembler performs well at <10X coverage and in *Eucalyptus pauciflora* genome assembly (Wang et al., 2020) where FLYE performs consistently well when >1 kb read lengths are subsampled when compared to >35kb read length subsamples. Iterative Racon polishing appears to have a beneficial effect (Figure 2). Across all read lengths a combined Medaka and Racon polishing strategy yields the most enhanced genome assembly (Figure 3).

Overall, this analysis highlights the advantages and shortcomings of various assemblers and provides use cases for each. SHASTA, when given higher coverage data, is an incredibly powerful assembler that runs quickly (Wick & Holt, 2019) and generates extremely accurate contigs. However, SHASTA is not robust in regards to data volume, as without sufficient read depth this assembler performs sub-optimally and fails to generate complete assemblies. It could still be useful at a lower depth of coverage for the purposes of complete and fast gene identification, particularly for large genomes. In comparison, CANU is the slowest running assembler, due to its extensive pre-assembly error correction and trimming steps. However, length can be prioritized over depth when using this assembler read and the incorporation of shorter read lengths may even result in sub-optimal results. The advantage of this assembler for more advanced users is the ability to modify parameters, however, this may not be appropriate for novices. FLYE is the most robust of the assemblers tested, with results across subsamples appearing consistent. This assembler may be an attractive tool for most data volumes and particularly for novice usage, as minimal parameter adjustments are required with the single caveat of a user-defined genome size.

### Hi-C Mapping performance across *K. excelsa* read length subsets

To assess the impact of initial assembly quality on Hi-C mapping performance, Hi-C data generated using the Phase Genomics kit was mapped to the initial assemblies generated across five *K. excelsa* read subsets. The contigs were further scaffolded using two commonly used software packages, AllHiC and SALSA2. AllHiC uses “pruning” and “optimization” steps to produce allele aware scaffolds, however, it requires *a priori* knowledge of the chromosome number. SALSA2 uses the ONT assembly graph in order to assess assembly accuracy prior to Hi-C scaffolding, and does not require karyotype information. Assessing assembly constructs from these two scaffolding software programs allowed not only a comparison of informed (AllHiC) and non-informed (SALSA2) strategies but also the performance of a software that corrects mis-assemblies prior to scaffolding to a software that scaffolds based on the input assembly alone. When comparing scaffolder performance, it was important to integrate quantitative metrics such as N50 and total length but also intrinsic karyotype information, as these quantitative metrics alone fail to determine over-assembly by the scaffolder and although give an indication of assembly accuracy they are unable to determine the scaffolders precision. This strategy has been integrated into the Vertebrate Genomes Project’s (Rhie McCarthy, et al., 2020) validation workflow in order to generate near error-free genome assemblies using Hi-C data.

Amongst all read length subsamples the resulting assemblies from the SALSA2 Hi-C scaffolding are more accurate than those produced by the AllHiC package. SALSA2 generates assemblies that are more complete and are of a total length closer to the expected genome size. These findings are further supported by Figure 5c and 5d that highlight suspicious scaffold length distributions constructed by AllHiC, a result of over-assembly. This over-assembly could be a consequence of the homozygous *K. excelsa* sample resulting in a reduced long range interaction signal (Zhang et al., 2019) and other more heterozygous genomes may indeed perform better.

However, AllHiC also fails to construct scaffolds from ONT assemblies with an abundance of short contigs, this shortcoming being highlighted in the failure of AllHiC to construct scaffolds for >5 kb and 10 kb read subsample assemblies produced by WTDBG2, which contain 10,317 and 10,043 contigs respectively. SALSA2 circumvents this issue by removing all contigs <1000 bp prior to HiC assembly.

Through shorter contig removal SALSA2 achieves scaffolds utilizing all WTDBG2 subsamples and overall performs better when longer reads are supplied, although the >30 kb read subsample SALSA2 scaffolding slightly underestimated the total length. As previously mentioned, all SHASTA generated assemblies, although gene complete, underestimate the total length of the genome and despite Hi-C scaffolding the assembly failed to gain coverage of the entire genome but retained gene completeness score. For instance, the SHASTA assembly constructed using the >30kb subsample and scaffolded with SALSA2’s gene completeness was 85%, which was the highest score achieved when compared to all other ONT assemblers but the total length was less than half the expected length. Similarly, SALSA2 performs optimally for both the CANU and FLYE assembly constructed using the >30kb subsample as it does with SHASTA.

From these analyses it is clear that the input assembly does substantially affect the accuracy of Hi-C scaffolding, as the issue of fragmentation found in ONT assemblies produced by WTDBG2 profoundly affected the scaffolding process, generating assemblies of low contiguity even after Hi-C scaffolding. Furthermore, the lack of genome length coverage of the initial SHASTA assemblies was not resolved with the addition of Hi-C data. Again, FLYE appears more robust than other assemblers to scaffolding software, however, AllHiC still produces a suspicious read length distribution suggesting over-assembly in individual subsamples. CANU assemblies perform sub-optimally using the AllHiC scaffolder but the high quality initial >30kb read subsample ONT assembly appears to remain the superior assembly post SALSA2 scaffolding.

### Pseudo-chromosome construction of the final *K. excelsa* genome

Scaffolding of the *K. excelsa* was carried out by AllHiC and SALSA2. Overall, this analysis highlighted the importance of ONT assembly accuracy prior to Hi-C scaffolding as errors found in these assemblies were unresolved by further scaffolding. Consistent with the assessment of the ONT contig assembly performance, both FLYE and CANU assemblies retained the greatest amount of the input data in scaffolds constructed when compared to WTDBG2, NECAT and SHASTA. This is highlighted by quantitative metrics such as high consensus quality (QV), a kmer completion scores peaking at 91% and 93% respectively and map back rates peaking at 84% for both. The more accurate CANU and FLYE based scaffolds appear to be caused by the higher accuracy of the underlying ONT assembly.

Interestingly, FLYE-based Hi-C assemblies appear to have a higher degree of gene completeness in comparison to CANU however they have slightly smaller total assembly lengths than expected from flow cytometry. FLYE initial assemblies also appear robust to different scaffolding strategies with similar results across both AllHiC and SALSA2 being achieved. In contrast, the CANU assemblies from SALSA2 appear to be superior. Focusing on N50, contiguity, gene completeness scores and total length alone lead to misleading conclusions about genome accuracy being drawn as AllHiC appeared more contiguous and had similar total lengths and gene completeness scores when compared to SALSA2 assemblies. Through the integration of scaffold length distributions as a quality metric, the accuracy of these assemblies could be more thoroughly evaluated as the karyotype shows all 14 chromosomes are of similar length (Hair & Beuzenberg, 1958) and this should be represented in the scaffolds produced after Hi-C data integration. Here, SALSA2 had a more realistic length distribution whilst AllHiC generated assemblies with length distributions inconsistent with the karyotype.

SHASTA, NECAT and WTDBG2 all failed to produce reliable scaffolded assemblies, suffering from collapsed genome lengths, low kmer completeness, and poor mapping back rates and therefore were not considered for further analyses.

### A pseudo-chromosome length near-complete genome assembly for Proteaceae

After initial scaffolding, quality metrics were assessed, and the initial FLYE ONT assemblies scaffolded with AllHiC and SALSA2 were selected for additional pseudo-chromosome construction. AllHiC produced high quality metrics with a gene completeness of 88%, N50 of 66.6 Mbp, kmer completeness of 91%, and a Hi-C read map-back rate of 80%. However, total length is lower than expected at 816 Mbp, and evidence of over-assembly was identified with 50% of the genome being placed on a single chromosome.

SALSA2 produced scaffolds also performed well, with a gene completeness score of 87%, a total length of 977Mbp, kmer completeness of 84%, a Hi-C read map-back rate of 84% and a chromosome length distribution in line with what is expected for this species. However, in this case contiguity suffered with an N50 of only 1.56 Mbp being obtained. Both assemblies were further validated for structure and orientation accuracy through comparison to macadamia linkage maps. This analysis highlighted the accuracy and orientation of both scaffolds sets. Despite SALSA2 being less contiguous the metrics suggested more accurate scaffolds, therefore this assembly was further scaffolded to increase contiguity. After additional scaffolding, contiguity was increased with an N50 of 114 Mbp, whilst retaining high quality a total length and kmer completion, however, gene completeness scores dropped by 8%. In order to assess this, the data was manually curated using Juicer and Juicebox. Here, a single mis-assembly was detected and manually curated resulting in a genome that is 91% kmer complete, 97.5Mbp in length, 90% gene complete (99% complete if considering Eukaryota dataset) and has an N50 of 114 Mbp.

This assembly is the first near-complete genome sequence for the Proteaceae clade and will provide invaluable information to the honey production industry in Aotearoa New Zealand, but also provides a reference for other Proteaceae in this clade.

## Conclusions

Our long read and Hi-C based assemblies of *K. excelsa* could potentially be useful as a benchmarking resource to be utilized regularly on release of new ONT assembly and Hi-C scaffolding tools. This will allow the continuous assessment of performance of new genomic packages across both read length and read depth. Furthermore, this could enhance the genomics community’s ability to make a more educated *de novo* genome assembly pipeline prior to assembly whilst also giving information on the data volume required. In the future it will be important that more assemblers, polishing mechanisms and Hi-C scaffolders are investigated and benchmarked. Finally, the *K. excelsa* assembly produced here will be used in the future to assess the genomic diversity of rewarewa across its natural range in Aotearoa New Zealand, in collaboration with Māori agribusinesses involved in the honey industry.

## Availability of Supporting Data and Materials

Permission from representatives of the indigenous people (Māori) was obtained for using the plant material used for this study. Further studies using this material, raw sequencing data and final genome assembly will require consent from the Māori iwi (tribe) who exercises guardianship for this material according to Aotearoa New Zealand’s Treaty of Waitangi and the international Nagoya protocol on the rights of indigenous peoples. Raw and analyzed data is available through the Manaaki Whenua Landcare Research data repository (https://doi.org/10.7931/paqg-kk20) with managed access. Access to this data will require permission from representatives of the Te Rarawa iwi (tribe).

## Availability of Source Code and Requirements

See Supplementary Material 1.

## Authors’ contributions

A.M., T.R.B. and D.C. conceived the study. J.M.P. collected samples and G.H. coordinated the engagement with Te Rarawa Anga Mua and the Komiti Kaitiaki for Warawara Ngahere. E.H. performed the laboratory work. A.M. analyzed and interpreted the data. S.S.C., C.W. and J.G. contributed to Hi-C and scaffolding analysis. A.M. and D.C. wrote the manuscript with input from all authors. D.C. and T.R.B. coordinated the project.

## Acknowledgements

The authors thank the Te Rarawa iwi (tribe) for supporting this research and approving publication of this manuscript. The authors also thank members of Genomics Aotearoa’s High Quality Genomes project, support from New Zealand eScience Infrastructure (NeSI) and Amali Thrimawithana (Plant & Food Research) and Abdul Batten (AgResearch) for useful comments on the manuscript. Leaf samples were collected by Peter Bellingham (Manaaki Whenua – Landcare Research) with field support from Mike White (Northland Regional Council). The Department of Conservation helped with collecting logistics and permits (CA-31615-OTH).

## Abbreviations

ONT: Oxford Nanopore Technology;
SMRT: (PacBio) Single Molecule Real-Time;
PE: (Illumina short) paired-end reads;
WGS: whole genome sequencing;
Hi-C: high-throughput chromosome conformation capture;
kb: kilobase pairs;
LAI: Long terminal repeat retrotransposons Assembly Index:
BUSCO: Benchmarking Universal Single-Copy Orthologs;
RX4: Racon polishing (4 rounds);
RX8: Racon polishing (8 rounds);
SR: sort read polishing using Pilon;
PH: purge_haplotigs software.

## Tables and figures captions

**Supplementary Material 1: Software packages versions and parameters.**

**Supplementary Material 2: Illustration of Oxford Nanopore Technologies (ONT) read quality for *Knightia excelsa* before and after rebase-calling.** A) Distribution of read lengths generated from the ONT library. B) Distribution of Q scores per read. C) Summary of Qscore (Y axis) compared to read length (X axis) with higher read counts colored green.

**Supplementary Material 3: Quality assessment of all *K. excelsa* assemblies for Hi-C scaffolding**

**Supplementary Material 4: Utilization of Macadamia linkage maps to validate the orientation and order of the *K. excelsa* Hi-C scaffolds.** A summary of the results of the mapping of 14 Macadamia linkage groups two the two FLYE/MEDAKA/PILON/ *K. excelsa* scaffolds generated using AllHiC and SALSA2. A) Whole genome alignment of the Left) FLYE-SALSA2 Hi-C assembly Right) FLYE-AllHiC Hi-C assembly against the Macadamia scaffold sets produced by ALLMAPS. B) Telomere positions of Right) Longest 21 scaffolds produced by FLYE-SALSA2 and B) 14 pseudo-chromosomes produced by FLYE-AllHiC. C) Unique markers found in the Macadamia linkage groups mapped to the FLYE-SALSA2 Hi-C assembly.

