## Supplementary figures and images for "An exploration of assembly strategies and quality metrics on the accuracy of the *Knightia excelsa* (rewarewa) genome"

### Supplementary_File_2

Supplementary Material 2:

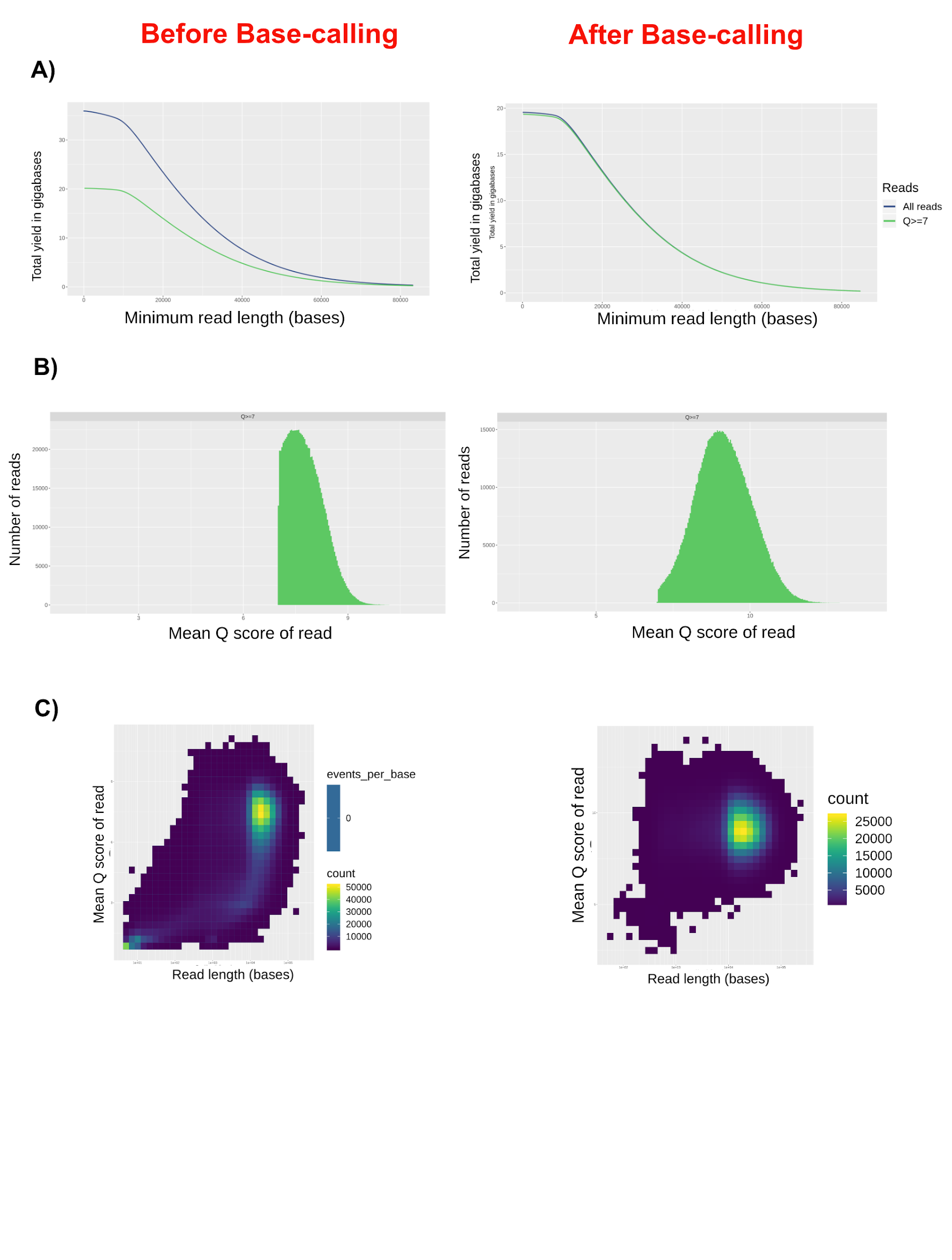

### Supplementary_File_4

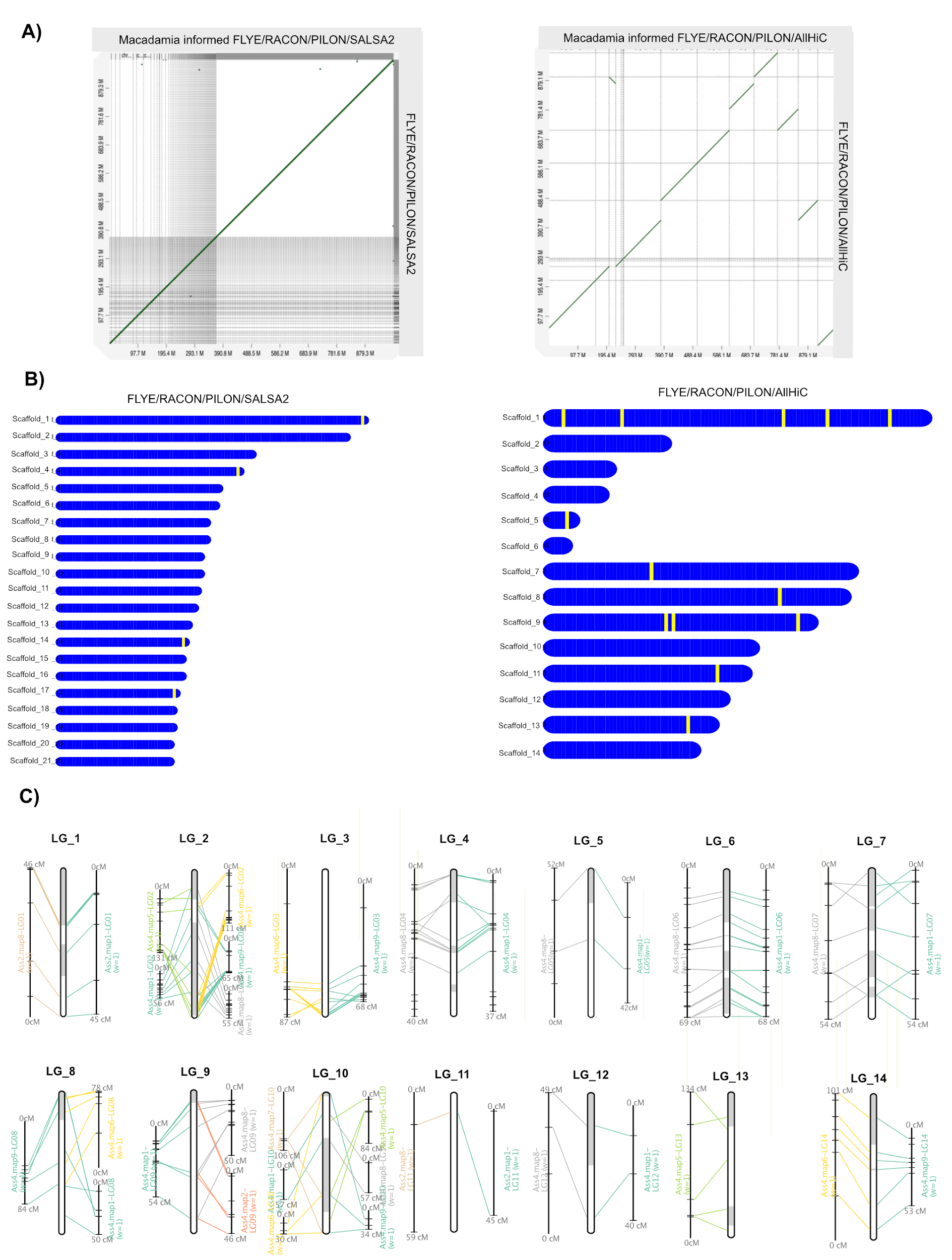
