## Supplementary_File_1 for "An exploration of assembly strategies and quality metrics on the accuracy of the *Knightia excelsa* (rewarewa) genome"

**Software package versions:**

**Guppy**, v2.2.3, **Porechop**v0.2.4 **BEDTools**v2.28.0, **SAMtools**v1.9, **Flye**v2.4.2, **Canu**v1.8, **WTDBG2**v2.5, **Pilon**v1.23, **Minimap2**v2.16, **BWA**v0.7.17, **Perl**v 5.28, **ALLHIC**v0.8.12, **BUSCO**v3.0.2, **QUAST**v5.0.2, **Shasta**v0.1.0, **NECAT** (<https://github.com/xiaochuanle/NECAT>), **Merqury**v1, **Meryl**v1, **networkx**v1.11, **MUMmer**v3.23, **quickmerge**v0.3, **Picard**v2.21, **Racon**v1.4, **medaka**v0.11.0, **PurgeHaplotigs**v1.1.1, **GenomeTools**v1.6.1, **LTR_retriever**v2.8.5, **BLAST**v2.9, **ALLMAPs**v1.0.6

**Computational Requirements and Parameters**

Parameters specified are those required for All_Data read subsample

**Assembly Workflows and Parameters:**

**Shasta Workflow**

Step_1: Fastq to fasta file conversion

python /nesi/nobackup/landcare02569/shasta/scripts/FastqToFasta.py FASTQ_FILE.fastq FASTA_FILE.fasta

Step_2: Shasta_Assembly

--cpus-per-task=30

--mem=2000G

--time=24:00:00

./shasta-Linux-0.1.0 --input FASTA_FILE.fasta

**Canu Assembly**

CANU v1.8 was compiled and run with jemalloc 5.2.1 (http://jemalloc.net/) to improve memory speed and access. The complete command is as follows, with input nanopore reads and output directory changing when appropriate for profiling with shorter reads.

LD_PRELOAD=`jemalloc-config --libdir`/libjemalloc.so.`jemalloc-config --revision` /software/canu-1.8/Linux-amd64/bin/canu -p Rewarewa -d Rewarewa genomeSize=1g -nanopore-raw rewarewa_rebasecalled.fastq >canu.log 2>canu.err

**Flye Workflow**

--cpus-per-task=10

--mem=1000G

--time=168:00:00

flye --nano-raw FASTQ_FILE.fastq -g 1g -o ASSEMBLY_DIRECTORY -t 10 -i 0

**Necat Workflow**

NOTE: For this assembler you need to generate multiple files: 1) Config.txt and 2) readlist.txt

Step_1: Config file construction

open nano and copy the following config_file format: Rewarewa config file used:

PROJECT=DIRECTORY_NAME

ONT_READ_LIST=readlist.txt

GENOME_SIZE=1000000000

THREADS=4

MIN_READ_LENGTH=3000

OVLP_FAST_OPTIONS="-n 500 -z 20 -b 2000 -e 0.5 -j 0 -u 1 -a 1000"

OVLP_SENSITIVE_OPTIONS="-n 500 -z 10 -e 0.5 -j 0 -u 1 -a 1000"

CNS_FAST_OPTIONS="-a 2000 -x 4 -y 12 -l 1000 -e 0.5 -p 0.8 -u 0"

CNS_SENSITIVE_OPTIONS="-a 2000 -x 4 -y 12 -l 1000 -e 0.5 -p 0.8 -u 0"

TRIM_OVLP_OPTIONS="-n 100 -z 10 -b 2000 -e 0.5 -j 1 -u 1 -a 400"

ASM_OVLP_OPTIONS="-n 100 -z 10 -b 2000 -e 0.5 -j 1 -u 0 -a 400"

NUM_ITER=2

CNS_OUTPUT_COVERAGE=45

CLEANUP=0

USE_GRID=false

GRID_NODE=0

FSA_OL_FILTER_OPTIONS="--max_overhang=-1 --min_identity=-1 --coverage=40"

FSA_ASSEMBLE_OPTIONS=""

FSA_CTG_BRIDGE_OPTIONS="--dump --read2ctg_min_identity=80 --read2ctg_min_coverage=4 --read2ctg_max_overhang=500 --read_min_length=5000 --ctg_min_length=1000 --read2ctg_min_aligned_length=5000 --select_branch=best"

Step_2: Readlist file construction

-- open nano and copy the path to your fastq files in it

nano readlist.txt

/path/to/your/fastq

Step_3: Assembly

Assembly Step 1:

--cpus-per-task=4

--mem=50G

--time=48:00:00

Prerequisites: Perl

perl necat.pl correct config.txt

Assembly Step 2:

--cpus-per-task=4

--mem=400G

--time=168:00:00

Prerequisites: Perl

perl necat.pl assemble config.txt

Assembly Step 3:

--cpus-per-task=4

--mem=400G

--time=168:00:00

perl necat.pl bridge config.txt

**Wtdbg2 Workflow**

Step_1:

--cpus-per-task=16

--mem=200G

--time=12:00:00

wtdbg2 -t 16 -g 1g -i input_reads.fastq -o assembly_output.fasta -x rs -p 19 -AS 2 -s 0.05 -L 5000

Step_2:

--cpus-per-task=16

--mem=200G

--time=42:00:00

/wtdbg2/wtpoa-cns -t 16 -i assembly_output.fasta.ctg.lay.gz -o assembly_output.fasta

**Purge Haplotigs Workflow**

Step_1:

--cpus-per-task=10

--mem=50G

--time=6:00:00

Prerequisite: SAMtools, minimap2, Perl, R, BEDTools

minimap2 -t 10 -ax map-ont pilon.fasta  long_reads.fastq > pilon.bam

samtools sort -@ 10 pilon.bam -o pilon_aligned.bam

purge_haplotigs readhist -t 10 -b pilon_aligned.bam -g pilon.fasta

**Note : Download .png generated file and interpret graph to fill in values for step 2, -l, -m, -h.

Step_2:

--cpus-per-task=20

--mem=50G

--time=12:00:00

Prerequisites: SAMtools, minimap2, Perl, R, BEDTools

purge_haplotigs contigcov -i pilon_aligned.bam.gencov -l **X** -m **X** -h **X** -o pilon_covstats.csv

purge_haplotigs purge -t 20 -g pilon.fasta -c pilon_covstats.csv –o Output_dir

**Assembly Polishing**

**Polishing Workflow**

Strategy 1) Minimap2 and Racon

--cpus-per-task=12

--mem=100G

--time=07:00:00

```
minimap2 -x ava-ont -t 12 "assembly.fasta" reads.fastq > minimap.paf
racon -t 12 reads.fastq minimap.paf  "assembly.fasta" > polished_assembly.fasta
```

Strategy 2) Medaka

--time=48:00:00

--mem=200G

--cpus-per-task=20

medaka_consensus -i bascalled.fastq -d input_assembly.fasta -m **model_required** -o output_dir –t 20

** Model used: r941_min_high_g303

Strategy 3) Pilon

--time 72:00:00

--mem 512G

--cpus-per-task 18

Prerequisites: Pilon, SAMtools, BWA, Java

bwa index input.fasta

bwa mem -t 18 "input.fasta" Shortread_1.fastq Shortread_2.fastq > input.sam

samtools view -u input.sam | samtools sort -o input.bam

samtools index input.bam

java -**Xmx160G** -jar $EBROOTPILON/pilon.jar --genome input.fasta --frags input.bam --outdir output_dirname --threads 18

**Hi-C Assembly**

**Phase Genomics Filtration Workflow**

Filtration and initial QC of HiC libraries for all HiC datasets were carried out using the following guidelines supplied by Phase Genomics (<https://phasegenomics.github.io/2019/09/19/hic-alignment-and-qc.html>).

**Hi-C Mapping Workflow**

Strategy 1: AllHiC

Step_1: Indexing

Prerequisites: SAMtools, BWA, Bedtools

bwa index -a bwtsw input_longread_assembly.fa

samtools faidx input_longread_assembly.fa

Step_2: Alignment

--cpus-per-task=24

--mem=50G

--time=48:00:00

Prerequisites: BWA

bwa aln -t 24 input_longread_assembly.fa Reads1.fastq > sample1_R1.sai

bwa aln -t 24 input_longread_assembly .fa Reads2.fastq > sample1_R2.sai

bwa sampe input_longread_assembly.fa sample1_R1.sai sample1_R2.sai Read1.fastq Read2.fastq > input_longread_assembly_aln.sam

OR

bwa mem -t 24 input_longread_assembly.fa Read1.fastq Read2.fastq > input_longread_assembly _aln.sam

Step_3: Filtration

--cpus-per-task=24

--mem=50G

--time=48:00:00

Prerequisites: Perl, SAMtools, BEDTools

/ALLHiC/scripts/PreprocessSAMs.pl input_longread_assembly_aln.sam input_longread_assembly.fa MBOI

Step_4: Bam Filtration

--cpus-per-task=24

--mem=50G

--time=48:00:00

Prerequisites: SAMtools

ALLHiC/scripts/filterBAM_forHiC.pl input_longread_assembly_aln.REduced.paired_only.bam input_longread_assembly_aln.clean.sam

samtools view -bt input_longread_assembly.fa.fai input_longread_assembly_aln.clean.sam > input_longread_assembly_aln.clean.bam

Step_5: Split reads in predefined clusters

--cpus-per-task=12

--mem=50G

--time=48:00:00

```
/ALLHiC/bin/ALLHiC_partition -b input_longread_assembly_aln.REduced.paired_only.bam -r input_longread_assembly.fa  -e GATC -k 14
**k=input your chromosome number
```

Step_6: CLM file generation and RE count

--cpus-per-task=24

--mem=50G

--time=48:00:00

/ALLHiC/bin/allhic extract input_longread_assembly_aln.REduced.paired_only.bam input_longread_assembly.fa --RE GATC

Step_7: Optimise

--cpus-per-task=12

--mem=50G

--time=24:00:00

/allhic optimize

Step_8: Assembly

Prerequisites: Perl

```
 perl ./ALLHiC_build input_longread_assembly.fa
for i in *.fasta; do perl ./../getFaLen.pl -i $i -o $i.len;done
grep 'counts_GATC' *.len > chrn.list
```

--cpus-per-task=18

--mem=50G

--time=14:00:00

Prerequisites: SAMtools, Python

./ALLHiC_plot input_longread_assembly_aln.REduced.paired_only.bam input_longread_assembly.groups.allhic.agp input_longread_assembly.groups.allhic.fasta.len 500k pdf

Strategy 2: Salsa2

Step_1:

--cpus-per-task=24

--mem=50G

--time=24:00:00

Prerequisite: SAMtools

samtools view -bt input_longread_assembly.fa.fai input_longread_assembly_aln.sam > input_longread_assembly.bam

--cpus-per-task=24

--mem=50G

--time=48:00:00

Prerequisites: BEDTools, SAMtools

bamToBed -i input_longread_assembly.bam > input_longread_assembly.bed

sort -k 4 input_longread_assembly.bed > tmp && mv tmp input_longread_assembly.bed

--cpus-per-task=24

--mem=50G

--time=48:00:00

Prerequisites: Networkx, Python

```
python /SALSA/run_pipeline.py -a input_longread_assembly.fa -l input_longread_assembly.fa.fai -b input_longread_assembly_sorted.bed -e GATC -o
 Input_longread_assembly -m yes
```

**Hi-C QC**

**Merqury Workflow**

Prerequisites: Meryl, Merqury

--time=48:00:00

--mem=100G

--cpus-per-task=10

Prerequisites: SAMtools, BEDTools, IGV,meryl

sh merqury.sh Database.k20.meryl/ HiC_Assembly.fasta outputdir

**Create meryl database prior to merqury

**LAI index**

Step1: Indexing

#SBATCH --cpus-per-task=4

#SBATCH --mem=20G

#SBATCH --time=02:00:00

Prerequistes: genometools, Samtools

samtools faidx scaffolds_FINAL.fasta

gt suffixerator -db scaffolds_FINAL.fasta -indexname hiC_LTR.fsa -tis -suf -lcp -des -ssp -sds -dna

Step2: LTR harvest (Obtain LTR regions)

--cpus-per-task=4

--mem=30G

--time=02:00:00

Prerequisites: genometools, LTRharvest

gt ltrharvest -index hiC_LTR.fsa > hiC_LTR.out

Step3: LTR_retriever (LAI index on LTRs)

--cpus-per-task=4

--mem=50G

--time=48:00:00

Prerequisites: BLAST, RepeatMasker, Perl

perl /LTR_retriever/LTR_retriever -genome scaffolds_FINAL.fasta -inharvest hiC_LTR.out > LTR_retriever_hiC.out
